# Global regulators facilitate adaptation to a phenotypic trade-off

**DOI:** 10.1101/2023.09.19.558433

**Authors:** Matthew Deyell, Vaitea Opuu, Andrew D. Griffiths, Sander J. Tans, Philippe Nghe

## Abstract

Organisms depend on their ability to balance multiple phenotypes by evolutionary adaptation. For instance, cellular growth and movement jointly enable critical processes including microbial colonization and cancer invasiveness. Growth and movement are known to be controlled by local regulators that target single operons, as well as by global regulators that impact hundreds of genes. However, how these different levels of regulation interplay during evolution is unclear. Using Escherichia coli growth and motility as a model system, we show that global regulators enable the adaptation of two phenotypes bound by a trade-off, where improvement in one causes deterioration in the other. We measured how CRISPR-mediated knockdowns of global and local transcription factors impact growth and motility in different environments. We find that local regulators mostly modulate motility, while global regulators jointly modulate growth and motility. Genetic perturbations display complex high order interactions between genes and environments. Nevertheless, gene perturbations display consistent patterns in the growth-motility space when grouped by their position in the regulatory hierarchy. These patterns constrain evolutionary scenarios, where local regulators are typically mutated first to optimize motility, then global regulators allow cells to adjust the trade-off between growth and motility. These findings overall highlight the role of pleiotropic regulators for coordinating phenotypic responses in complex environments.

## Introduction

The fitness of organisms depends on multiple phenotypes, but it is not always possible to optimize all of them simultaneously due to trade-offs [1]. A trade-off means that improving one trait may come at the cost of another. A key example is the trade-off between growth and motility, which has been demonstrated in bacteria and cancer cells [2-6]. Disseminated tumour cells are highly mobile, which promotes metastasis, but have a slow-cycling state that makes radiation therapy and chemotherapies poorly effective [2]. In bacteria, the trade-off between mobility and growth underlies the maintenance of population diversity through niche formation [6]. In the presence of a trade-off, evolutionary adaptation requires to adjust several phenotypes, either leading to specialization or to a balance between traits [7].

Here, we aim to test the hypothesis that global regulators play a role in coordinating multiple traits during evolution. Global regulators are defined as transcriptional factors which bind hundreds of operons, in contrast to local regulators, which are dedicated to one or a few [8]. Global regulators are primarily characterized at the molecular level, based on the knowledge of their binding sites [9]. Due to the number and diversity of operons they regulate, global regulators are expected to alter several phenotypes. For instance, CRP regulates the expression of secondary catabolites in response to cAMP [10]and is implicated in biofilm formation [11]. Other global regulators such as HNS and Fis (so-called Nucleoic Associated Proteins or NAPs) alter the compaction of the bacterial genome [12] depending on the growth phase [13] and participate to the stress response [14] and biofilm formation [15]. However, so far, studies of global regulators are focused on molecular mechanisms and consider a single phenotype [16]. Thus, how global regulators may coordinate multiple phenotypes in an evolutionary context is an open question.

To investigate the role global regulators during adaptation to a phenotypic trade-off, we used a model system in which local and global regulators regulate growth and swimming in E. coli. CRISPR interference allowed us to knock-down (denoted KD) transcription factors in combination, and hence study their genetic interactions. We chose 2 local (FlhDC and FliZ), 2 intermediate (mcaS and CsgD), and 3 global regulators (CRP, Fis, HNS), known to regulate each other [9] (Figure 1A). FliZ and FlhDC directly control expression of the flagella operon essential for swimming mobility [17]. CsgD controls expression of the Curli motility pathway and adhesion pili while repressing swimming [18]. CsgD is itself controlled by the small inhibitory RNA mcaS [19]. These mobility-regulating transcription factors themselves are upstream regulated by global transcription factors CRP, Fis, and HNS [9], described above.

**Figure 1.**
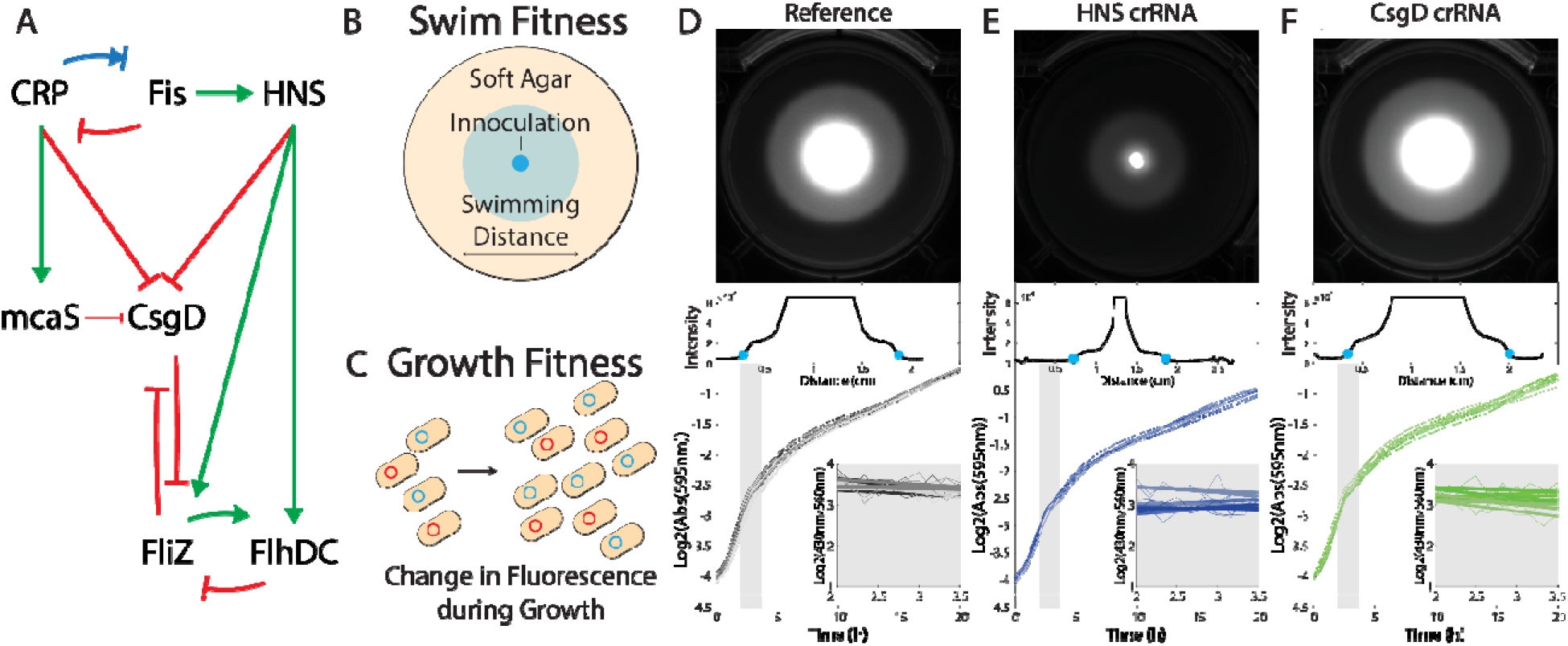
Quantification of swimming and growth. (A) Regulatory relationships between the selected panel of transcriptio factors. Each of these transcription factors was targeted with CRISPRi perturbation, either alone or in combination, an compared with a reference strain (RS) harbouring a non-targeting crRNA. (B) Swimming distance was determined through inoculation in the centre of soft agar plates and measurement of the diameter up to the swimming front. (C) Growth performance was determined as the change in fluorescence over time of the perturbation strain (cyan fluorescent) grown i competition with a reference strain (red fluorescent). (D) Raw data for the RS. (E) Raw data from the HNS perturbation, whic results in a decrease in swimming (top) but an increase in relative growth rate (bottom, slope). (F) Raw data from the CsgD perturbation, which results in a slight increase in swimming (top) but a slight decrease in relative growth rate (bottom).

Our approach allowed us to analyse the coupled variations of growth and swimming in response to KD of genes with different positions in the regulatory hierarchy (upstream, intermediate, or downstream), alone or in combination. We examined the genetic interactions (epistasis) between pairs of regulators and found that epistasis differs between the two phenotypes and is modulated by the environment, demonstrating higher-order interactions. Despite this complexity, the phenotypic effects of genetic perturbations follow typical patterns when grouping genes by their position in the regulatory hierarchy: mutations in local regulators mostly improve swimming while global regulators adjust the balance between growth and swimming. Based on these measurements, we simulated evolution in variable environments where fitness depends on both phenotypes. We found that genetic changes in local regulators generally precede those in global regulators, where the latter navigate between optimal phenotype combinations.

## Results

### Characterizing relationships between genetic perturbations, phenotypes, and environments

We used CRISPR interference (CRISPRi) to generate single and double knockdown (KD) perturbations of 7 transcription factors (CRP, Fis, HNS, mcaS, CsgD, FliZ, FlhDC). Cells were transformed with vectors expressing an inducible catalytically deactivated Cas9 (dCas9), mCerulean to fluorescently tag the cells, and a CRISPR RNA (crRNA) that targets the transcription start sequence of the relevant transcription factor (see Methods). CRISPRi represses gene expression by using the dCas9 as a programmable transcription factor [20]. We then quantified the swimming and growth phenotypes for this E. coli strain library (Figure 1), and the unperturbed reference strain (RS).

Swimming was quantified using a migration assay in 0.3% soft agar plates. Mobile bacteria form concentric bands while swimming in soft agar plates (Figure 1BDEF). These bands arise due to local gradients of nutrients formed by cell consumption [21, 22]. We quantified swimming as the diameter of the first swimming wave after 16 hours in LB media or 24 hours in M63 media. Growth was quantified using a competition assay against an unperturbed RS in microwell plates with constant shaking (Figure 1CDEF). Growth was determined based on three signals: optical density, mCerulean fluorescence (emission 430 nm) of the query strain, and mCherry fluorescence (emission 560 nm) of the reference stain. The relative growth rate is computed as the slope of the logarithm of the ratio between fluorescence signals and is restricted to the exponential phase as controlled using optical density at 595 nm (OD). This growth rate is normalized using the OD and fluorescent measurements to remove contributions due to differences between constructs and fluorescent markers, including maturation times, degradation rates, and expression costs (see Supplementary Text).

Swimming and growth were both measured over 9 replicates, consisting of 3 biological replicates (different days and initial colonies), each comprising 3 technical replicates (same day, different wells or plates). The swimming and growth values were taken as the median of all 9 measurements during the exponential growth phase and were highly reproducible (Supplementary Figure 1, Supplementary Figure 2). In a given medium, differences between swimming and growth values across strains were significant, confirming the effectiveness of the KDs on these phenotypes (Supplementary Figure 1, Supplementary Figure 2).

### Regulatory mutants are bound by a growth-motility trade-off

To understand how genetic perturbations impact phenotypes, we examine the distribution of phenotypes within the 2D space consisting of the growth rate (x-axis) and swimming distance (y-axis) for single and double KD strains (28 KD strains in total) in the 3 environments (Figure 2). Single and double KDs cause mild to strong effects on both phenotypes, with growth rate differences ranging from -0.17 h ^1^ to 1.09 h^-1^, and swimming diameter ranging from 0.4 cm to 3.4 cm after 16 hours (see Supplementary Table 1 for RS values). Growth is observed to increase or decrease due to the genetic perturbations while swimming distance only decreases.

**Figure 2.**
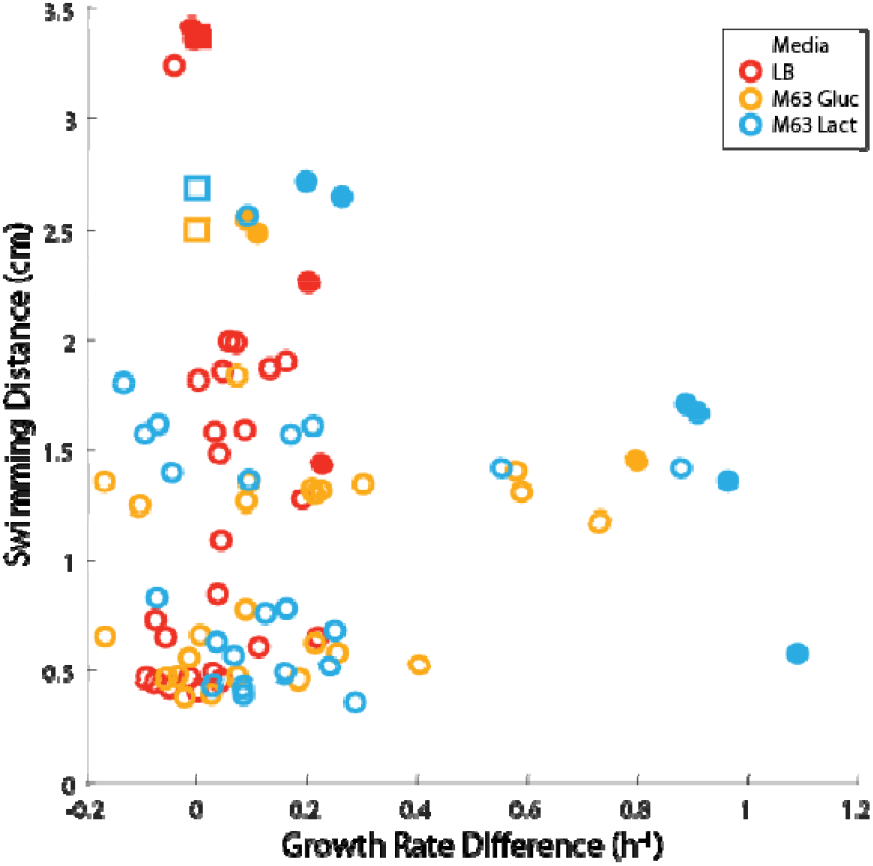
Growth-swimming phenotype space displays a Pareto front. The distance that each strain can swim in 16 hours plotted against the growth rate difference with the RS (squares) for each KD strain (circles). Solid symbols are Pareto optim strains.

Environments have contrasting effects on the phenotypes. On the one hand, they alter the range of accessible growth values, which varies less for rich media (Figure 2). Indeed, the largest relative changes in growth rates are observed in the poorest medium M63 Lactose (1.24-fold change), followed by M63 Glucose (0.97) and finally LB (0.32). On the other hand, the range of accessible swimming performances is quite similar across environments. Mainly, the best performing strain in LB swims 0.35-fold and 0.25-fold further than the best strains in M63 with glucose and lactose media, respectively.

Moreover, the distribution of genetic variants in the phenotypic space of Figure 2 displays a trade-off: mutants with a growth rate higher than the unperturbed RS (squares in Figure 2) swim comparatively less. This trade-off between growth and swimming was already characterized in experimental adaptive evolution experiments and mutants [34, 35]. Such trade-offs are analysed using the concept of Pareto optimality [7], where organisms that cannot be outcompeted on all functions simultaneously are said to be Pareto optimal. In Figure 2, Pareto optimal strains are such that no other strain grows more without swimming less. Pareto optimal strains localize on a line called the Pareto front [1]. Respectively 4, 6, and 3 Pareto-optimal strains are observed in LB, M63 Glucose and M63 Lactose (solid symbols in Figure 2).

### The coupling between phenotypes depends on gene hierarchy

To analyse the relationship between regulatory network structure and phenotypic effects, we first examined whether changes in growth and swimming were specifically associated to the genes being perturbed (Figure 3). Each panel of Figure 3 reports the phenotypic effects of a given knockdown (KD) across all genetic backgrounds and environments. A vector along the x-axis means the KD affects growth only, while a vector along the y-axis means the KD affects swimming only. Diagonal vectors indicate pleiotropy, in other words, a coupling between growth and swimming. If the coupling is positive, growth and swimming improve both (top-right quadrant) or deteriorate both (bottom left quadrant). If the coupling is negative, improved growth comes at the expense of deteriorated swimming (bottom-right quadrant), or vice versa, growth deteriorates while swimming improves (top-left quadrant).

**Figure 3.**
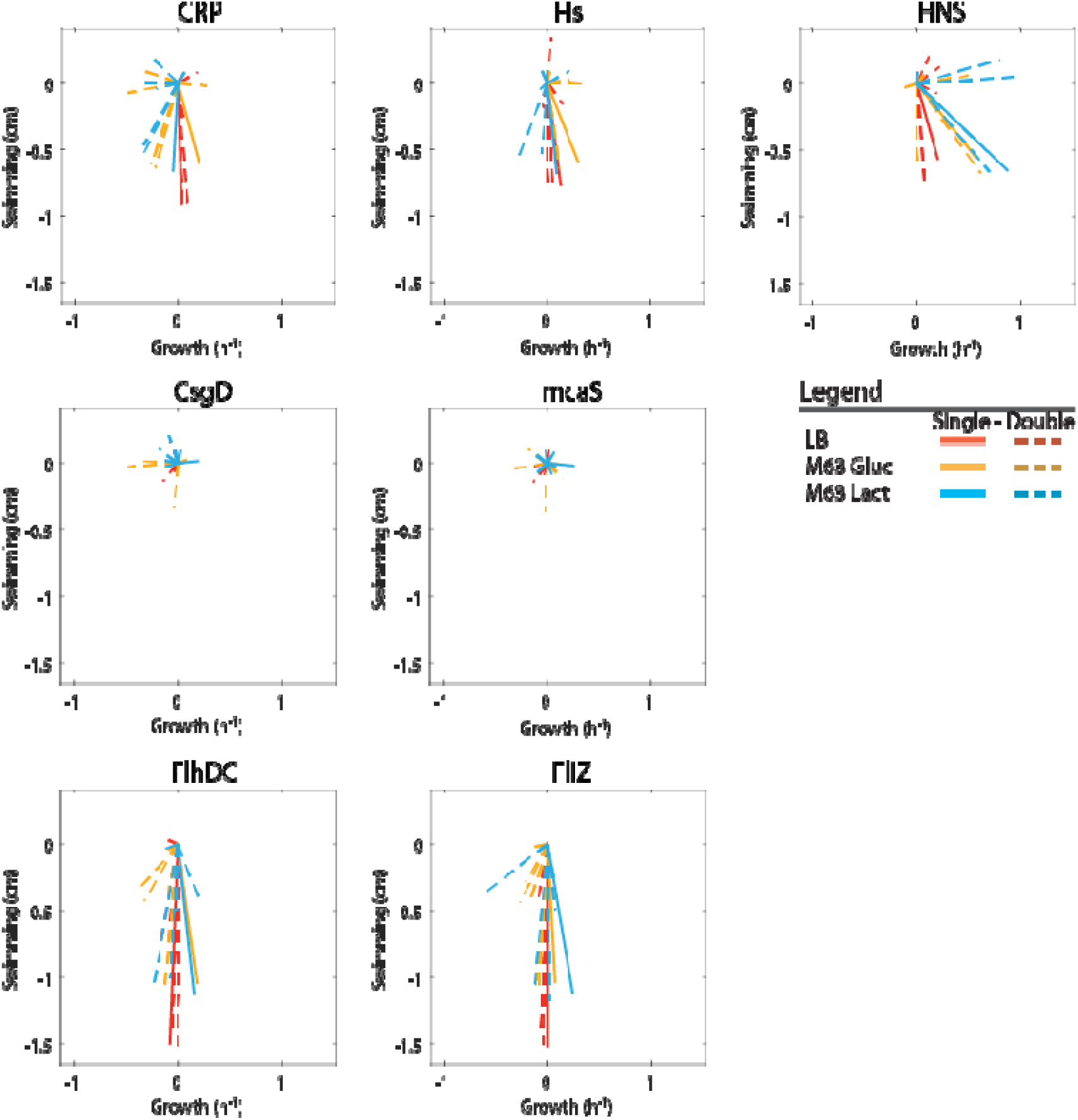
Impact of regulatory structure on the magnitude and direction of phenotypic effects. The vectors represent the phenotypic difference in the growth-swimming space for each given KD, where the strain without the KD is at the origi irrespective of its genotype.

KDs yield a varying range of couplings between phenotypes depending on the gene being perturbed. CRP, Fis, and HNS show the most diverse responses to their KD, where different genetic backgrounds and environments lead to changes in growth exclusively, swimming exclusively, or both (Figure 3). Perturbations of mcaS and csgD also had varied effects but of low magnitude as the majority are not statistically significant (p>0.05 Welch t-test n = 9). In contrast, FliZ and FlhDC had strong effects on motility regardless of environment, and showed a mild positive or negative impact on growth.

The 3 groups of transcription factors described above correspond to different positions within the regulatory network hierarchy (Figure 1). CRP, Fis, and HNS are upstream global regulators. Together, they accounted for 69% of perturbations that show negative growth-motility coupling (vectors that point top-left or bottom-right in Figure 3), mostly caused by HNS (n = 16). Such couplings are of particular interest as manifestations of the growth-motility trade-off. The regulators with the mildest effects, mcaS and csgD, occur in an intermediate position in the network hierarchy. Finally, FliZ and FlhDC had the strongest impact on swimming but little impact on growth. This is consistent with their position as downstream transcription factors dedicated to flagella regulation. FliZ and FlhDC accounted for 60% (6/10) of the few cases where swimming and growth both phenotypes decreased (7% of all perturbations).

This analysis indicates that downstream local transcription factors mostly modulate swimming, as may be expected, but upstream global transcription factors couple growth and swimming in a variety of ways, where swimming and growth may be decoupled, both increase, both decrease, or vary in opposite directions. However, it remains unclear at this stage how the diversity of effects caused by perturbations in global regulators leads to the distribution phenotypes observed in Figure 2.

### Genetic interactions are high-order but consistent with regulatory hierarchy

To understand the influence of regulatory interactions on phenotypic effects, we first quantified the epistasis between each pair of transcription factors [23], which quantifies the departure from additive effects. Epistasis may take various forms including magnitude, masking, or sign epistasis (Figure 4A). Magnitude epistasis conserves the sign of the fitness effect across backgrounds but not the amplitude of the effects. We denote here the extreme form of magnitude epistasis as ‘masking epistasis’, in which a genetic perturbation has no further effect if another perturbation has already occurred (in accordance with the classical definition of phenotypic epistasis). Sign epistasis refers to the case where a mutation causes a fitness increase in one background but a decrease in another background. When sign epistasis applies to both genes being perturbed, it is referred to as reciprocal sign epistasis.

**Figure 4.**
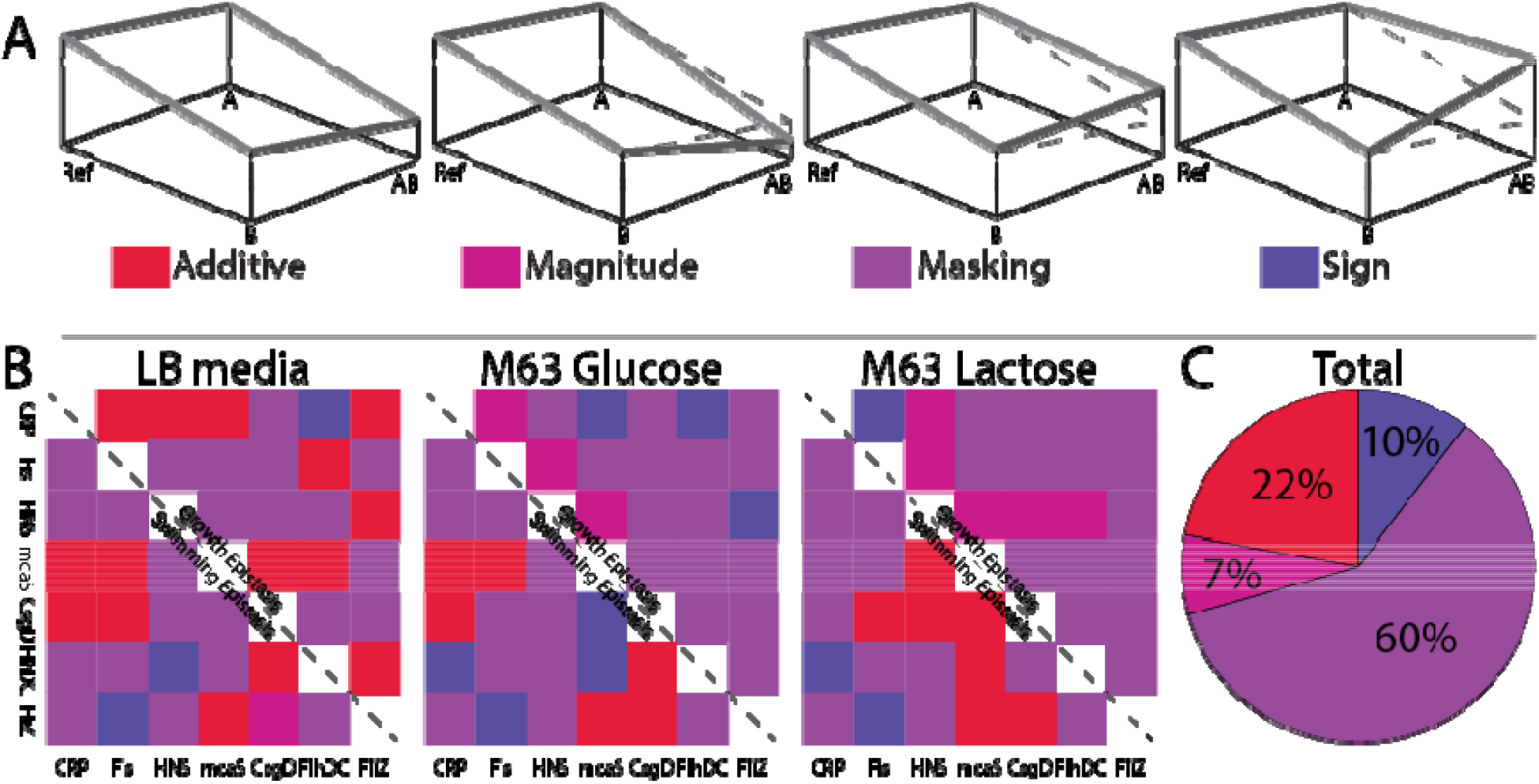
Epistasis between regulatory gene knock-downs. (**A**) Additive case (no epistasis, parallel edges): mutation A has the same effect irrespective of the background (RS or mutant B), and mutation B has the same effect irrespective of the background (wild-type [ref] or mutant A). Magnitude epistasis: the effect of mutation A is increased or decreased by the presence of mutation B, but the direction of the effect is the same (same for mutation B in the presence or not of mutation A). Maskin epistasis: mutation A has no effect in the presence of mutation B. Sign epistasis: the direction of the effect of mutation A is changed by the presence of mutation B. (**B**) Calculated Epistasis for each fitness metric in each growth media. Epistasis was calculated as described in figure 4. Most observed epistasis is masking, followed by additive. Total number of conditions tested is 126, n = 9. (C) Epistasis statistics observed across all perturbation pairs and media.

In total, we categorized 126 epistasis interactions, resulting from 21 gene pairs for 2 phenotypes across 3 environments (Figure 4BC, detailed in Supplementary Figure 3, 4 and 5). Epistasis is pervasive with only 22% of gene pairs showing additive effects across all environments and for both phenotypes (Figure 4BC). It has been suggested that epistasis may be determined by the relative position of genes within the upstream-downstream hierarchy of regulatory cascade [24]. However, this is not the case in our dataset, as epistasis differs in the majority of the cases phenotypes for a given pair of genes, as seen by comparing the colour of the squares positioned symmetrically across the diagonal in Figure 4B: over a total of 21 gene pairs, epistasis differs in 17 cases in LB, 13 cases in M63 Glucose, and 13 n M63 Lactose. This discrepancy with former observations may result from the regulatory network being more entangled than a mere gene regulatory cascade, with feed-forward loops and mutual feedbacks [25]. Genes display higher order effect with the environment. Indeed, for a given gene pair and a given phenotype, epistasis differs in at least one of the 3 environments in 27 over 42 cases, consistent with other studies [26].

The above analysis reveals the complexity of genetic interactions but does yet indicate any obvious relationship between genes, phenotypes and environments. However, a careful examination of the combined effects of gene KDs in the joint space of growth and swimming (Supplementary Figures 3, 4 and 5) points to typical patterns when grouping genes as upstream (global regulators CRP, Fis, HNS), intermediates (mcaS, CsgD), and downstream (local regulators FlhDC, FliZ). For instance, interactions between upstream and other upstream or intermediate genes (first 3 columns, top 5 rows in Supplementary Figures 3, 4 and 5) lead to triangular patterns. Interactions between upstream and downstream genes (first 3 columns, bottom 2 rows) tend to yield larger perturbations in both swimming and growth as compared to interactions between other ensembles. Interactions between downstream genes (bottom 2 rows) and other downstream intermediate genes (last 4 columns) are dominated by a strong decrease in swimming with little impact on growth.

The insight gained from the hierarchical grouping of gene invites us to re-examine the overall set of perturbations in the phenotypic space (Figure 5). We coloured each edge by which of the 3 gene groups was affected by the genetic perturbation: upstream global (yellow), intermediate (purple), or downstream local (blue) regulators. In this representation, the overall structure of gene perturbation effects appears consistent across environments (Figure 5A-C) even though particular epistasis for a specific pair of KDs may not be (Figure 4 and Supplementary Figure 4). In Figure 5, we observe that significant changes in swimming (vertical edges) are mainly attributed to local (blue) or global (yellow) perturbations. Significant changes in growth (horizontal edges) mostly occur via global regulators (yellow) irrespective of the swimming value, and intermediate regulators (purple) for strains that swim moderately well. Positively coupled changes in phenotypes (oblique edges with a positive slope) occur mostly via local regulators, starting from low swimming strains. Negatively coupled changes in phenotypes (oblique edges with a negative slope) occur almost only via global regulators, and trade swimming for growth.

**Figure 5.**
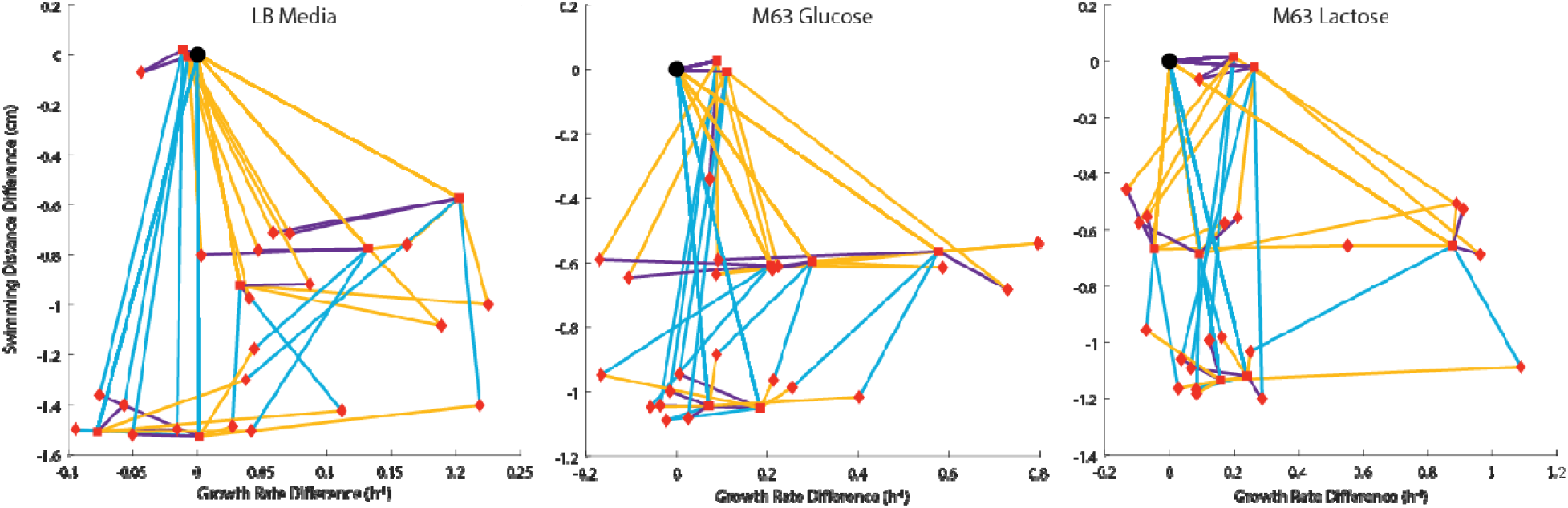
Relationship between regulatory network structure and response in phenotype space. The trajectories for eac perturbation are coloured by the group of the transcriptional regulator: Global (yellow), Intermediate (Purple), or Local (blue). The reference strain is represented as a black circle, single perturbations are represented as a square, and double perturbations are represented as a diamond.

Overall, local regulator perturbations mostly connect strains poor at both phenotypes to strains nearby the Pareto front, which are either good swimmers or moderate swimmers with large growth. In contrast, global regulator perturbations have strong effect on growth in the low swimming region, but trade swimming for growth in the Pareto front region. This suggests that, during evolution, local regulators perturbations are required to reach the Pareto front, but global regulators ultimately allow fine-tuning along the Pareto front.

### Evolution under complex selection pressures involves local then global regulators

To investigate the role of local and global regulators during adaptation, we computed evolutionary trajectories in variable environments where fitness depends on the two phenotypes. We consider here the regime where environments vary on timescales typically longer than that require for mutations to fixate. In the following, for simplicity, we call ‘mutation’ any genotypic perturbation, which in our experiment corresponds to a loss of function (KD) or a gain of function (KD taken in reverse, or knock-up KU).

We consider fitness functions where the relative importance of growth and swimming is controlled by a weighting parameter λ: When λ=0, only swimming determines fitness; When λ=1, only growth determines fitness. When λ=0, we defined the positive contribution of swimming to fitness using a sigmoidal function of the swimming distance with parameters s_0_ and k to (Figure 6A). The parameter s_0_ is the typical swimming distance difference with the reference strain allowing survival, which we set to - 1 cm, the middle of the range of the measured values (a higher or lower s_0_ would make swimming differences irrelevant to selection). The parameter k determines the sigmoid sharpness, where a very small k makes the response flat (no impact of swimming) and a very large k tends to a step function (threshold selection). When λ=1, fitness is taken as the differential growth rate normalized over the range of growth in a given medium. Intermediate values of λ interpolate between swimming and growth fitness functions. We explored the regime of strong selection for evolutionary trajectories within the 6 landscapes generated by KD pairs comprising a global regulator (CRP or Fis or HNS) and a local regulator (FlhDC or FliZ). For each landscape, we used the experimentally measured phenotypic values of the 4 genotypes and simulated evolutionary trajectories in an environment that varies randomly between the 3 media, as exemplified for the pair HNS-FlhDC in Figure 6B. The starting genotype is taken randomly for each realization of the simulation, which allows us to record statistics of the mutational paths over all possible adaptation scenario. Populations rapidly reach an equilibrium between mutation and selection after each environmental change, where they display small fluctuations around average proportions between genotypes (Figure 6B).

**Figure 6.**
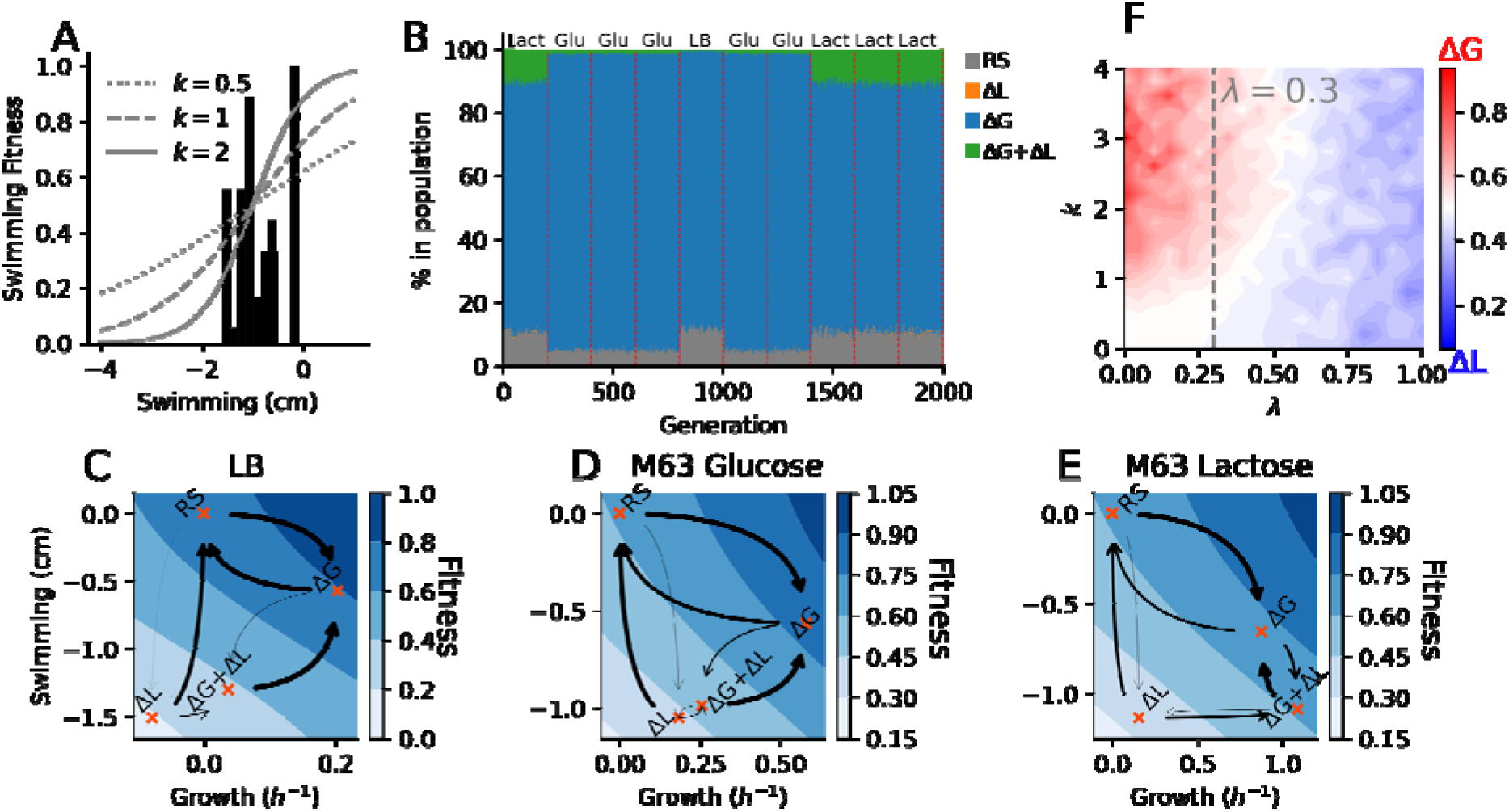
Evolution in variable environments. A) Distribution of swimming distance (histogram) and examples of swimming-to-fitness relationships, computed as a sigmoid of the swimming distance parameterized by a steepness parameter k, here centred on s_0_ = -1 cm. B) Example of evolutionary trajectory between the 4 genotypes (RS, ΔL, ΔG, ΔL+ΔG) generated by HNS and FlhDC knock-outs in a variable environment randomly alternating between LB (‘LB’), M63 Glucose (‘glu’) and M63 Lactose (‘lact’) every 200 generations. C-E) Mutational trajectories for the HNS-FlhDC in the three environments. Arrow thickness is proportional to the rate of transition during evolution. The fitness landscape is depicted by a gradient of blue. F) Frequency of mutations by type across the 6 combinations of mutations involving a global regulator (CRP, Fis, HNS) and a local regulator (FlhDC; FliZ), following the protocol of panel B. G) Proportion of local and global mutations as a function of k and λ (s_0_ = -1 cm).

We first focused on regimes where the contributions of growth and swimming to fitness are well-balanced. This is realized around λ=0.3, where fitness range is similar on the x and y axes (equivalently, isofitness lines are oblique, see Figure 6CDE for the pair FlhDC-HNS, and Supplementary Figure 6B for all pairs). In the following, ΔL denotes the strain with a local regulator KD, ΔG the strain with a global regulator KD, and ΔL+ΔG the double KD. Figure 6CDE and Supplementary Figure 5 depict the mutational landscapes in different environments and for different choices of local-global regulator pairs, with arrows representing mutations between genotypes. The thickness of an arrows is proportional to the probability to mutate from a genotype to another.

We observe typical trajectories that are reproduced between all mutational landscapes. For instance, in panels CDE of Figure 6, thick arrows depart from the ΔL and ΔL+ΔG genotypes, but those genotypes have weak incoming arrows. This indicates that ΔL and ΔL+ΔG are populated only transiently and tend to evolve irreversibly toward other genotypes. In contrast, the RS and ΔG genotypes have strong incoming and outgoing arrows, indicating that evolution can occur back and forth between those two genotypes located on the Pareto front (Figure 6CDE). The dynamics just described in the case of the HNS-FlhDC pair is also observed in 14 of the 18 landscapes (Supplementary Figure 5B), with first a knock-up of the local regulator (from ΔL to RS or from ΔL+ΔG to ΔG, depending on the starting genotype), then an exchange between the RS and the ΔG genotypes via the global regulator. Exceptions occur in lactose, with the above mentioned FlhDC-HNS pair as it also includes the ΔL+ΔG genotype on the Pareto front (Figure 6E), and the pairs FlhDC-CRP, FliZ-CRP, and FliZ-Fis, which display collapsed trajectories dominated by an exchange between ΔL and RS (Supplementary Figure 6B). Nevertheless, genotype exchange via the global regulator is the dominant mode of long-term evolution in variable environments, as confirmed on average over all simulated evolutionary trajectories (Figure 6F, λ=0.3), as well as for each pair considered separately (Supplementary Figure 6, λ=0.3).

In regimes where fitness is dominated either by growth or by swimming, the local or global regulator may be the ultimate target of adaptation. For instance, a k close to 0 or λ close to 1 corresponds to fitness being dominated by growth (Supplementary Figure 5A). Mutations in local regulators dominate (Figure 6F, Supplementary Figure 7), due to genotypes with maximal growth being in large majority ΔG, with ΔL+ΔG coming second, both exchanging via a local regulator mutation (12 over 18 pairs over the 3 media, Supplementary Table 2, Supplementary Figure 6). For k not too small and λ close to 0, fitness is dominated by swimming (Supplementary Figure 6C). Mutations occur mostly in the global regulator (Figure 6F, Supplementary Figure 7) due to the RS swimming the furthest and the ΔG coming second (Supplementary Figure 6C).

## Discussion

Transcription factors CRP, HNS and Fis are global regulators, which regulate hundreds of operons in bacteria. They have been detected from large scale mapping of binding sites and are known to be implicated in various phenotype, such as metabolism, biofilm formation and motility. Furthermore, global transcription factors are frequently modified when bacteria undergo directed evolution in unforeseen complex environments [27]. Overall, this hints at their role in jointly optimizing multiple phenotypes. However, no study had consistently examined the impact of perturbations in global regulators in a context where multiple phenotypes determine fitness.

Here, using genetic perturbations in E coli, we have shown that global regulators help adjust the relative contribution of growth and swimming, when those two phenotypes reach nearly optimal values. This is significant because growth and swimming are in a trade-off, and selective pressures may favour one or the other. Additionally, our analysis of evolutionary scenarios show that global regulators may not act alone: they must interplay with local regulators, which are required in early evolutionary steps to improve the swimming phenotype [28].

These findings invite us to reconsider possible explanations for the origin of global regulators. One explanation is that they are regulatory ‘hubs’ which result from a random process of network wiring [29]. Network hubs can result from the rule of preferential attachment, where transcription factors targeting many genes have more chance to regulate newly appeared genes than those dedicated to a single operon (e.g. following gene duplication-divergence events) [30]. However, it is unclear whether such a network wiring process make sense in the context of selection.

A contrasting explanation of global regulators is functional. It posits that co-regulation of operons by global and local regulators reflects cellular decision-making. For instance, alternative sugar operons are triggered only in the absence of glucose when CRP is activated. This is the case of the lac operon, which additionally requires the allosteric suppression of LacI repression in the presence of lactose. The underlying rationale is a prioritization between carbon sources depending on their quality to ensure optimal metabolic efficiency [31]. However, even though HNS and Fis may potentially follow a similar logic, for example triggering biofilm formation as a function of the growth phase, this has not been demonstrated. Global regulation may also come with an evolutionary cost due to their highly pleiotropic character, as mutating them can cause a large number of deleterious effects [32].

The present study highlights an additional ingredient, central to complex adaptations: the existence of phenotypic trade-offs. Adaptation in the presence of trade-offs requires the simultaneous and coordinated optimization of multiple phenotypes. Evolution then becomes dependent on genetic changes that couple variations in multiple phenotypes [33]. In this context, accessible evolutionary trajectories require some degree of pleiotropy, as we observed when only global regulators can adjust two optimal phenotypes. Pleiotropy may be a necessary evil to adapt to phenotypic trade-offs, thus illustrating how network complexity may find its roots in functional complexity [34].

## Methods

### Creation of vectors

pCRRNA mCherry and pCRRNA EGFP were made by Gibson Assembly. PCR was performed on pMD019 (pGFP) and pMD024 (pmCherry) with primers oMD546 and oMD547, and on pCRRNA with primers oMD545 and oMD548. PCR products were purified with PCR clean up kit from Macherey-Nagel. Equal molar concentrations of PCR product were mixed (one from either pMD019 or pMD024 and one from pCRRNA) and 5 µL of mix was added to 15 µL of Gibson Master mix to create pCRRNA Green and pCRRNA Red respectively. Golden Gate Assembly was used to replace the TrrnB terminator from pMD019 and pMD024 with B0014 as the TrrnB terminator contained 2 BsaI sites. PCR was done on pCRRNA Green and pCRRNA Red with primers oMD609 and oMD610, and on pCKDL with primers oMD607 and oMD608. PCR products where joined with Golden Gate Assembly using BsmBI enzyme to create pCRRNA EGFP and pCRRNA mCherry. pCRRNA mCerulean was created by Gibson Assembly. PCR was performed on pMD027 (pCerulean) with primers oMD704 and oMD705, and on pCRRNA mCherry with primers oMD706 and oMD707). PCR products were purified with PCR clean up kit from Macherey-Nagel. Equal molar concentrations of PCR product were mixed and 5 µL of mix was added to 15 µL of Gibson Master mix. All vectors were sequenced by GATC Biotech prior to use.

### Growth Conditions of Cultures

The host strain for all pCRRNA vectors is MG1655 with the pdCas9 vector from Stanley Qi (provided by Lun Cui and David Bikard). Glycerol stocks of each culture were streaked onto individual Lysogeny Broth (LB) agar plates containing 34 µg/mL of Chloramphenicol and 50 µg/mL of Kanamycin. Single colonies were inoculated into 2 mL of selected media (either LB or M63 supplemented with 0.4% Glucose or Lactose) containing 34 µg/mL Chloramphenicol and 50 µg/mL Kanamycin. Cultures were placed in a 37°C incubator for either overnight for 16 hours for LB cultures or for 24 hours for M63 cultures.

### Growth Competition Assays

Assays were performed by diluting 220 µL pCRRNA mCherry NT pre-culture into 22 mL of selected media containing 34 µg/mL Chloramphenicol, 50 µg/mL Kanamycin, and 25 ng/mL anhydrotetracycline. A Greiner, 96 Well, PS, F-Bottom, µCLEAR, Black microplate was filled with 198 µL of diluted culture per well. For each pCRRNA EGFP knockdown vector, 2 µL of pre-culture was inoculated into 3 individual wells. For the Non-targeting Control strain ‘RS, 2 µL of pre-culture was inoculated into 15 individual wells. A volume of 60 µL of mineral oil was added to each well of the microplate. The absorbance at 595 nm, as well as fluorescence at 480/510 nm and 580/610 nm (excitation/emission) were recorded every 10 minutes for 20 hours with a SpectraMax i3x. Microplates were incubated at 37°C and shook for 90 seconds before and after each measurement. The absolute growth rate of the reference strain was determined by the change in the log2(OD595nm) / time of the exponential growth phase (50mins to 170mins in LB, 50mins to 230mins in M63). To determine the fitness measurement of each knockdown, the ratio of the green and red fluorescence of the strain was divided by the median ratio of the green and red fluorescence of the 12 additional wells for the non-targeting control. Fitness measurements for each knockdown were made for LB media, and M63 media containing 0.4% of either glucose or lactose.

### Swimming Fitness Assays

For each knockdown, 10 µL of pre-culture was spotted into the middle of 15 mL of selected media soft agar (0.3%) plates. Plates where then incubated at 37°C for either 16 hours for LB plates or 24 hours for M63 plates. Plates where then imaged using a USB Camera and python viewer. An image of a non-inoculated plate was subtracted from each image. The fitness was determined by the ratio of the swimming area diameter of each knockdown strain to that of the non-targeting control strain.

### Evolutionary simulation

We defined our fitness function as: *F* = *λg* + (*1 − λ*)*Sig*(*k* × (*s* − *s*_*0*_)), where g and s are the growth and swim phenotypic values, respectively, λ controls the relative importance of both phenotypes, 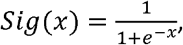, *S*_0_ is the inflection point of the sigmoid function, and k determines its steepness. For each KD landscape, we have four values of measured growth and four values of measured swim for each pair of genes, allowing us to simulate their dynamics.

We simulated the gene dynamics using the following genetic algorithm with mutation and bottleneck, looped through steps 2-4 for N generations:

#### 1. Initialization

We initialized a population of P individuals, each randomly assigned to one of the four possible genotypes: RS, local KD, global KD, double KD.

#### 2. Mutation

2% of the population was mutated (strong selection regime). For this subset, we randomly chose one of the two genes and knocked its state either up or down.

#### 3. Evaluation

We computed the fitness for each individual in the population.

#### 4. Selection

We implemented a selection process by subsampling the population based on the probability of each individual i: 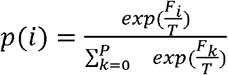, where T represents a fictitious temperature that allows us to adjust the strength of the selection.

For each KD pair, the environment was randomly changed every 200 generations, for a total of 2000 generations at T=7.

### Codes

The code is written in Python version 3.10.9 and has been run on a Linux machine with 20 cores 12th Gen Intel(R) Core(TM) i7. A simulation for a population size of 2000 and 2000 generations takes less than a minute. The code to perform the simulations and reproduce the figures are provided at https://doi.org/10.5281/zenodo.10201172, which are in the format of notebooks for the analysis, and Python scripts for the routines.

The routine evo_loop_env_change from the src/evo_four_states.py script performs the evolutionary simulation. It requires the measured phenotypic parameters located in the data directory. Then, the user can choose the population size, maximum number of generations, the frequency at which the environment is changed, and the mutation rate in the population.

In output of the simulation routine, the populations as well as the mutations accepted are recorded for each generation in two variables. In these variables, genotypes are encoded in the following way RS = (- 1, -1); Delta Local = (1, -1); Delta Global (−1, 1); and double knock down = (1, 1).

## Acknowledgments

The authors acknowledge the French Agence Nationale pour la Recherche ANR GeWiEpi (ANR-18-CE35-0005-01) and Institut Pierre-Gilles de Gennes Investissements d’avenir program (ANR-10-EQPX-34).

