## Supplemental Information and Figures for "Global regulators facilitate adaptation to a phenotypic trade-off"

**Supplementary Information and Figures**

Fitness differences between a mutant and the reference strain (respectively denoted with superscripts ‘m’ and ‘w’) are measured as the slope of the function:

$$loglog \left( \frac{F_{r}^{m}}{F_{b}^{w}} \right)\left( \frac{F_{r}^{w}}{F_{b}^{w}} \right)$$

where $F$ is the fluorescence measured over time with indices ‘r’ and ‘b’ for mCherry and CFP fluorescent reporter strains. To show that this measurement is appropriate, let us first evaluate how a fluorescent intensity $F_{i}$ depends on time. First, given the fluorescence intensity $a_{i}$ per protein of type ‘$i$’ and a maturation time $\tau_{i}$, fluorescence is delayed compared to the number of accumulated fluorescent proteins $N_{i}$, so that:

$F_{i}(t)=a_{i}N_{i}(t-\tau_{i})$	(1)

Second, during steady exponential growth at rate $\lambda_{i}$, proteins are produced at a rate proportional to the total mass of the cell, and considering the degradation rate $d_{i}$:

$$\frac{dN_{i}}{dt}= N_{i}^{0}e^{\lambda_{i}t}-d_{i}N_{i}$$

The solution to the above equation is:

$$N_{i}\left( t \right)=N_{i}^{0}e^{\lambda_{i}t}\frac{1-e^{-(\lambda_{i}+d_{i})t}}{\lambda_{i}+d_{i}}$$

The term $e^{-(\lambda_{i}+d_{i})t}$ fades exponentially compared to 1, with a characteristic time shorter than a cell cycle by definition of $\lambda_{i}$. Consequently, after a few cycles, $N_{i}$ is very well approximated as

$$N_{i}\left( t \right)\approx\frac{N_{i}^{0}e^{\lambda_{i}t}}{\lambda_{i}+d_{i}}$$

In combination with (1), this gives:$F_{i}\left( t \right)=\frac{a_{i}N_{i}^{0}}{\lambda_{i}+d_{i}}e^{\lambda_{i}(t-\tau_{i})}$

The ratio between the intensity of any two fluorescent signals is:

$$loglog \left( \frac{F_{2}}{F_{1}} \right) =\left[ \frac{a_{2}N_{2}^{0}}{a_{1}N_{1}^{0}}\frac{\lambda_{1}+d_{1}}{\lambda_{2}+d_{2}}+\lambda_{1}\tau_{1}-\lambda_{2}\tau_{2} \right]+\left( \lambda_{2}-\lambda_{1} \right)t$$

Therefore, the slope $\left( \lambda_{2}-\lambda_{1} \right)$ as a function of time corresponds to the fitness difference, any dependence in maturation time, initial amount, and degradation rate being accounted for by the bracketed constant term. Still, a fluorescent strain may be affected by a growth cost $c_{i}$ specifically associated with the fluorescent protein being expressed, so that it measured fitness $\lambda_{i}$ differs from its fitness $\mu_{i}$ in the absence of the reporter:

$$\lambda_{i}=\mu_{i}-c_{i}$$

The contribution of $c_{i}$ is removed when computing the slope of $loglog \left( \frac{F_{r}^{m}}{F_{b}^{w}} \right)\left( \frac{F_{b}^{w}}{F_{r}^{w}} \right)$, as, given the above, this slope equals the fitness difference independently of the construct and initial conditions:

$\left( \lambda_{r}^{m}-\lambda_{b}^{w} \right)-\left( \lambda_{b}^{w}-\lambda_{r}^{w} \right)=\left( \mu^{m}-c_{r}^{m}-\mu^{w}+c_{b}^{w} \right)-\left( \mu^{w}-c_{b}^{w}-\mu^{w}+c_{r}^{w} \right)=\mu^{m}-\mu^{w}$.

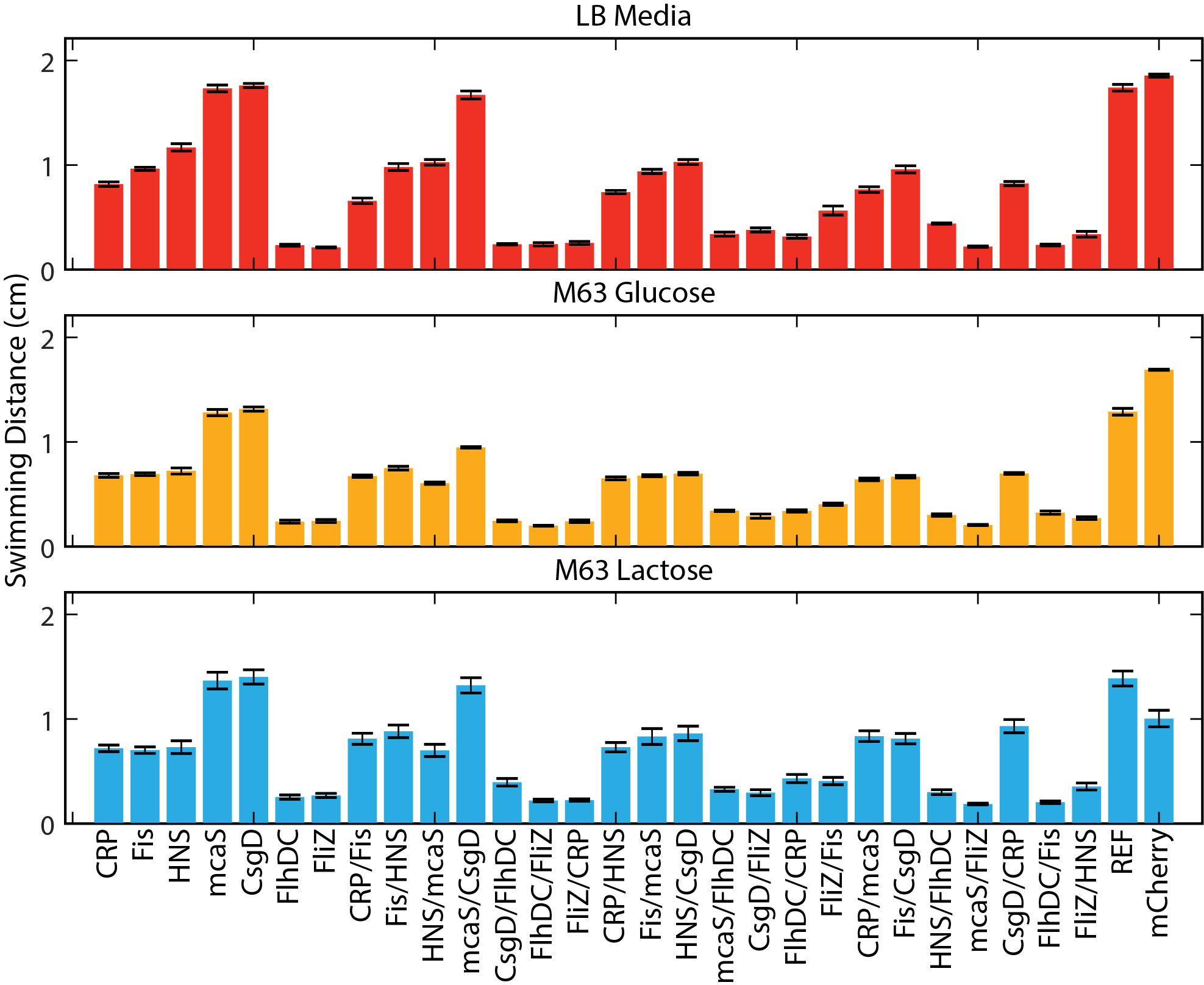

**Supplementary Figure 1**: *Mean values (n = 9) for swimming distance in soft agar after 16 hours from each CRISPRi perturbation grown in competition with non-targeting reference. Error bars represent SEM.*

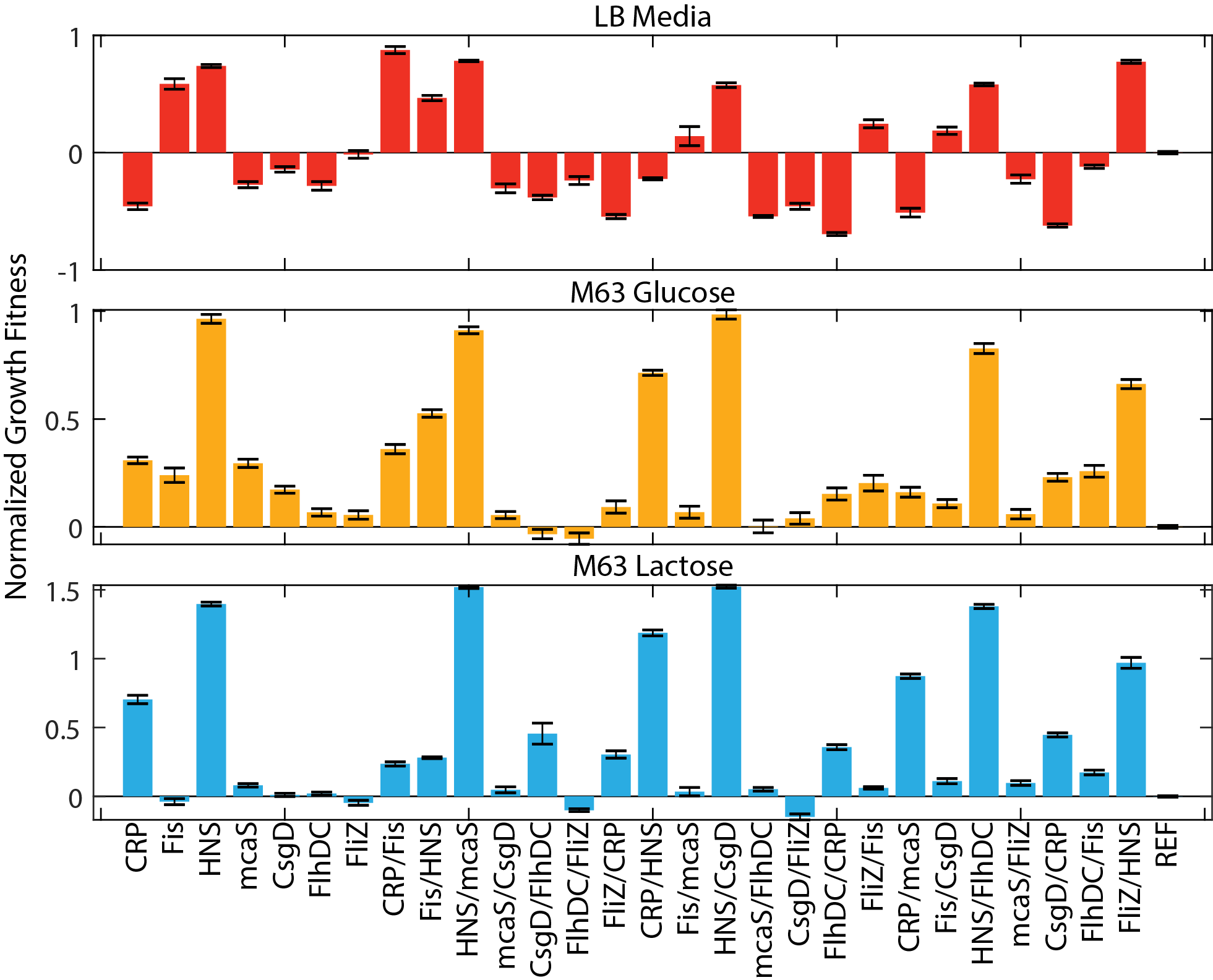

**Supplementary Figure 2***: Mean values (n = 9) for normalized growth fitness from each CRISPRi perturbation grown in competition with non-targeting reference for 20 hours. Error bars represent SEM.*

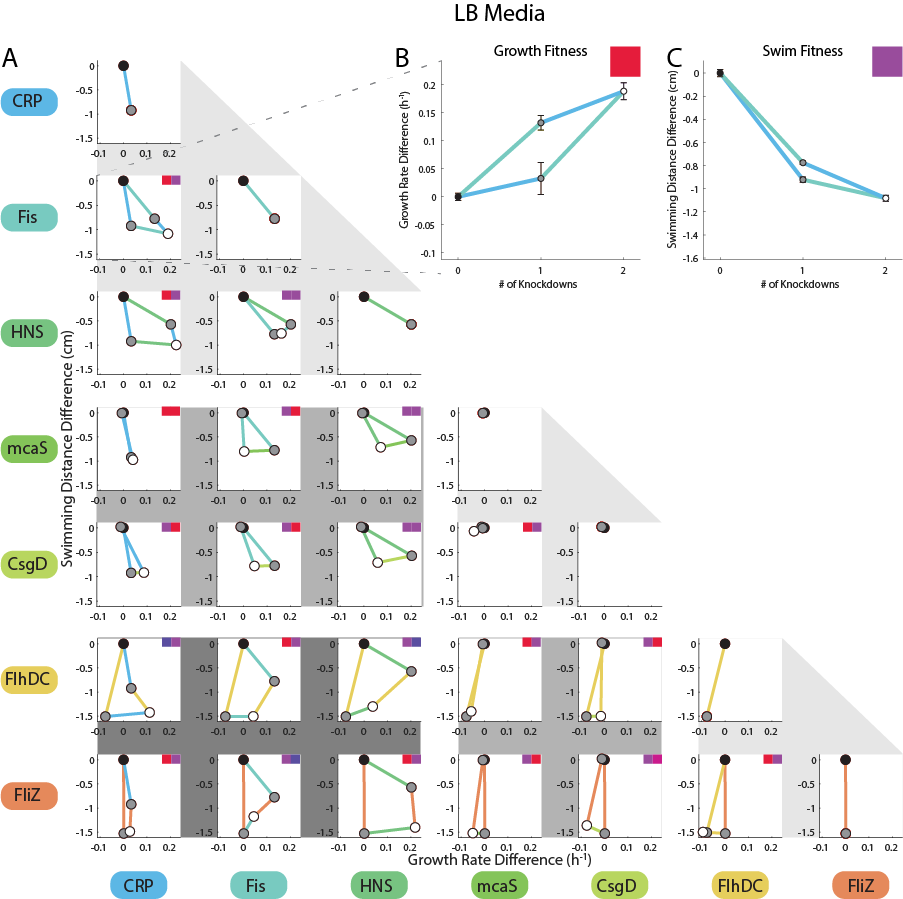

**Supplementary Figure 3**: *Each subpanel displays the growth and swimming values of a KD pair in LB media. The black dot is the unperturbed reference strain, the grey dots the single KDs, and the white dot the double KD. Edges are KD responses coloured in accordance with the gene lists on the left and bottom. Each subpanel results from 2 epistatic patterns: one for growth and one for swimming, as exemplified for the CRP-Fis pair in the right and left inserts, respectively. The pair of small squares in each subpanel indicates the epistasis type for growth (left square) and swimming (right square) following the colour code of panel A in Figure 3 of the main text. The background grey shadings highlight sets of interactions consistent with regulatory hierarchy: light grey are within group interactions (local-local, intermediate-intermediate, global-global), average grey are interactions between intermediate and other groups, dark grey are global-local interactions.*

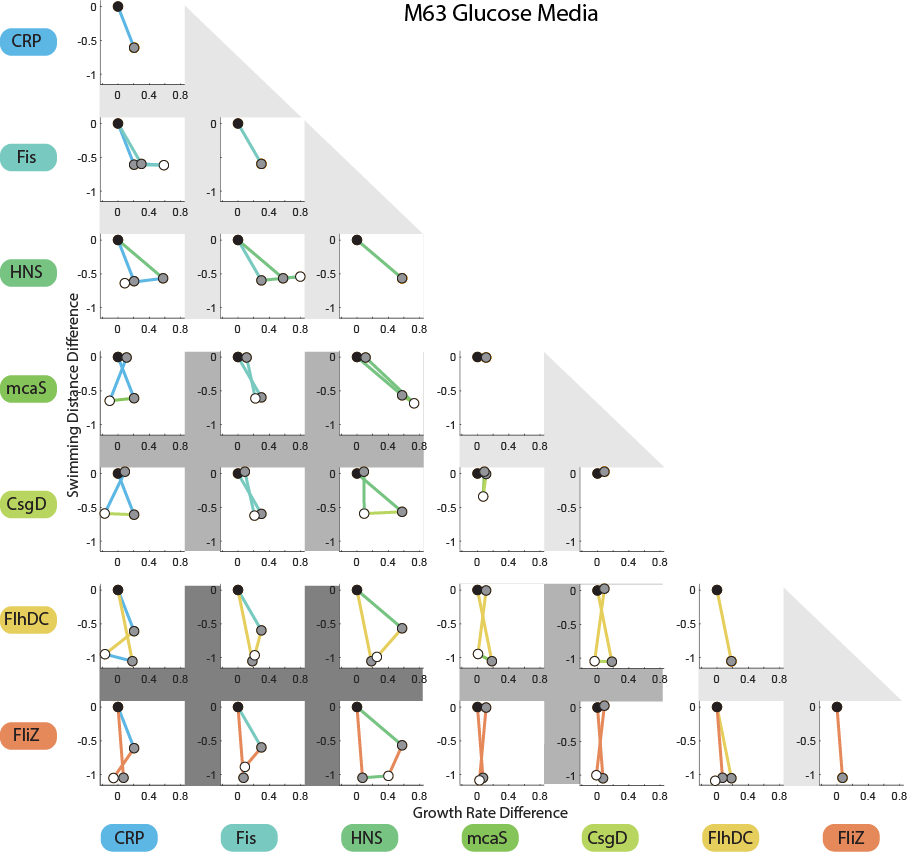

**Supplementary Figure 4**: *Each subpanel displays the growth and swimming values of a KD pair in M63 Glucose. The black dot is the unperturbed reference strain, the grey dots the single KDs, and the white dot the double KD. Edges are KD responses coloured in accordance with the gene lists on the left and bottom. Each subpanel results from 2 epistatic patterns: one for growth and one for swimming, as exemplified for the CRP-Fis pair in the right and left inserts, respectively. The pair of small squares in each subpanel indicates the epistasis type for growth (left square) and swimming (right square) following the colour code of panel A in Figure 3 of the main text. The background grey shadings highlight sets of interactions consistent with regulatory hierarchy: light grey are within group interactions (local-local, intermediate-intermediate, global-global), average grey are interactions between intermediate and other groups, dark grey are global-local interactions.*

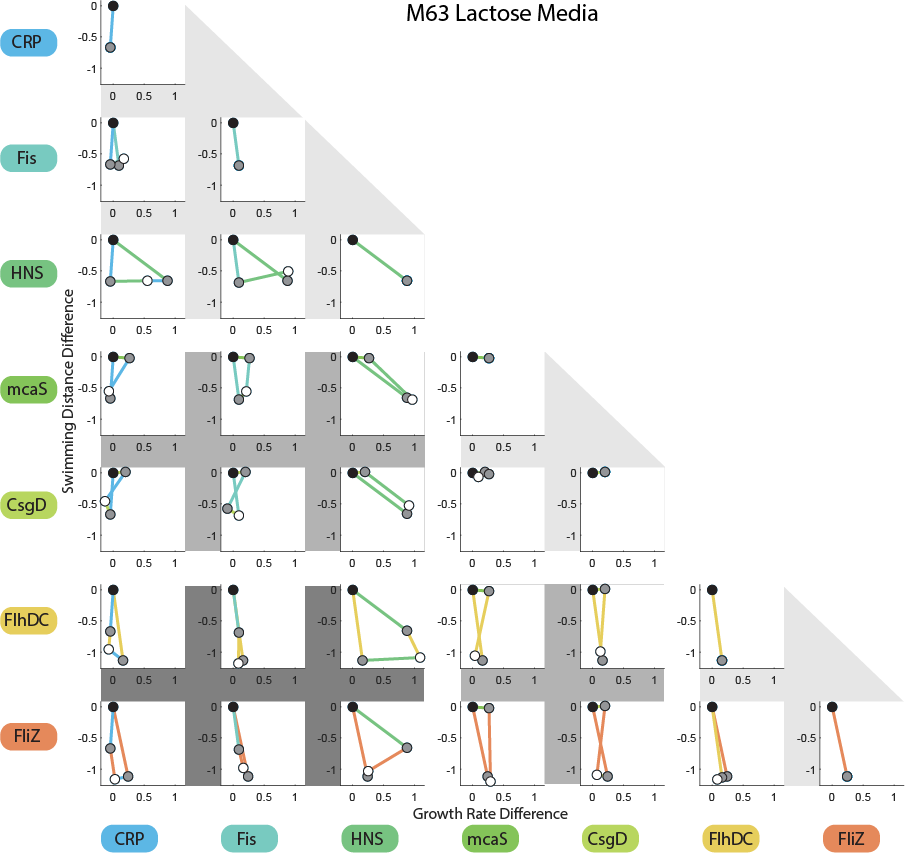

**Supplementary Figure 5**: *Each subpanel displays the growth and swimming values of a KD pair in M63 Lactose. The black dot is the unperturbed reference strain, the grey dots the single KDs, and the white dot the double KD. Edges are KD responses coloured in accordance with the gene lists on the left and bottom. Each subpanel results from 2 epistatic patterns: one for growth and one for swimming, as exemplified for the CRP-Fis pair in the right and left inserts, respectively. The pair of small squares in each subpanel indicates the epistasis type for growth (left square) and swimming (right square) following the colour code of panel A in Figure 3 of the main text. The background grey shadings highlight sets of interactions consistent with regulatory hierarchy: light grey are within group interactions (local-local, intermediate-intermediate, global-global), average grey are interactions between intermediate and other groups, dark grey are global-local interactions.*

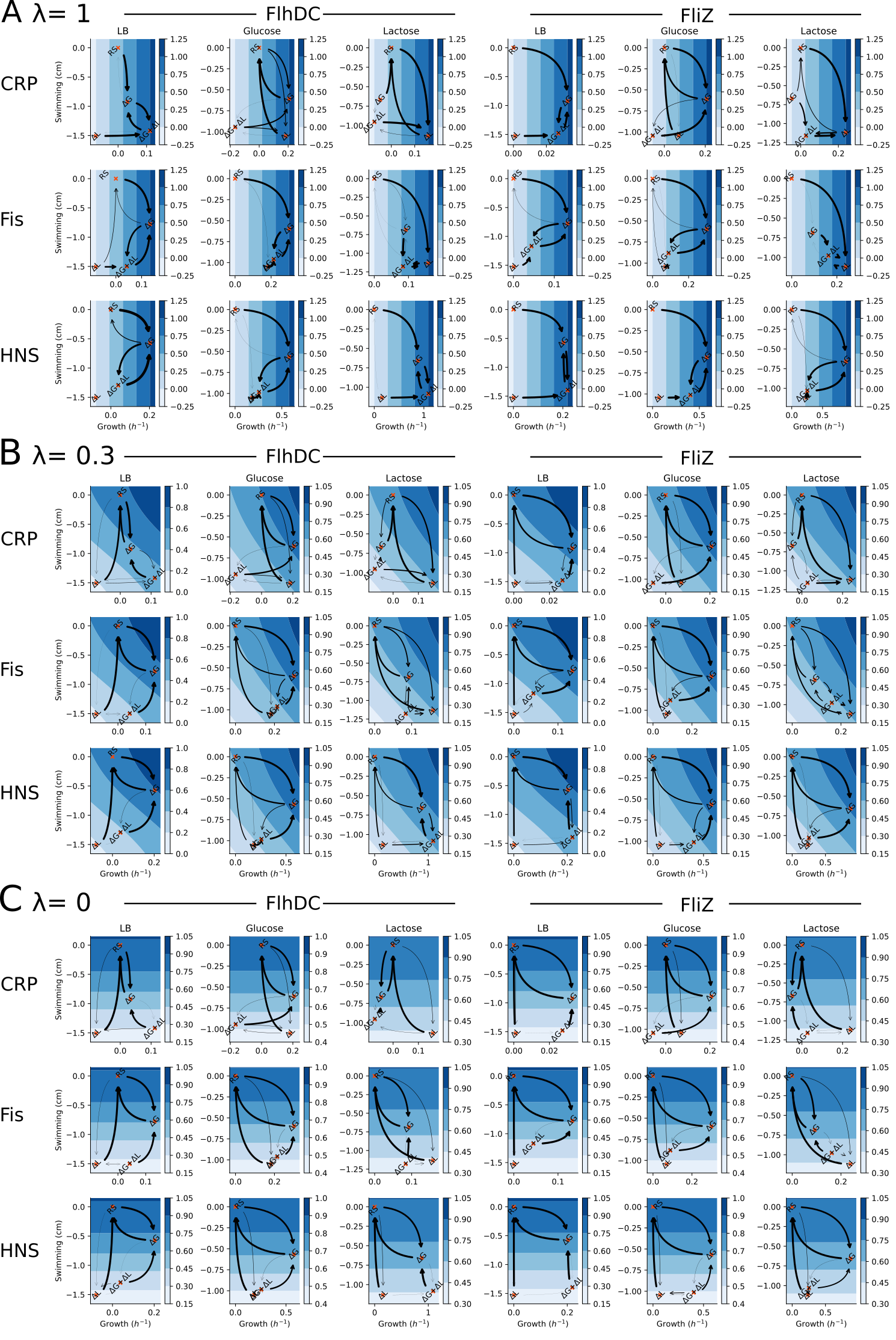

**Supplementary Figure 6**: *Mutational trajectories for each of the 6 KD pairs combining a global regulator (CRP, Fis, HNS) and a local regulator (FlhDC, FliZ) in variable environments randomly switching between LB, M63 glucose, and M63 lactose. Arrow thickness is proportional to the proportion of the mutation from the strain. The y-axes are the swimming difference with the RS in cm unit, the x-axis are the growth rate difference in h^-1^.* ***(A)*** *λ=1, only growth contributes to fitness;* ***(B)*** *λ=0.3, growth and swimming equally contribute to fitness;* ***(C)*** *λ=0, only swimming contributes to fitness.*

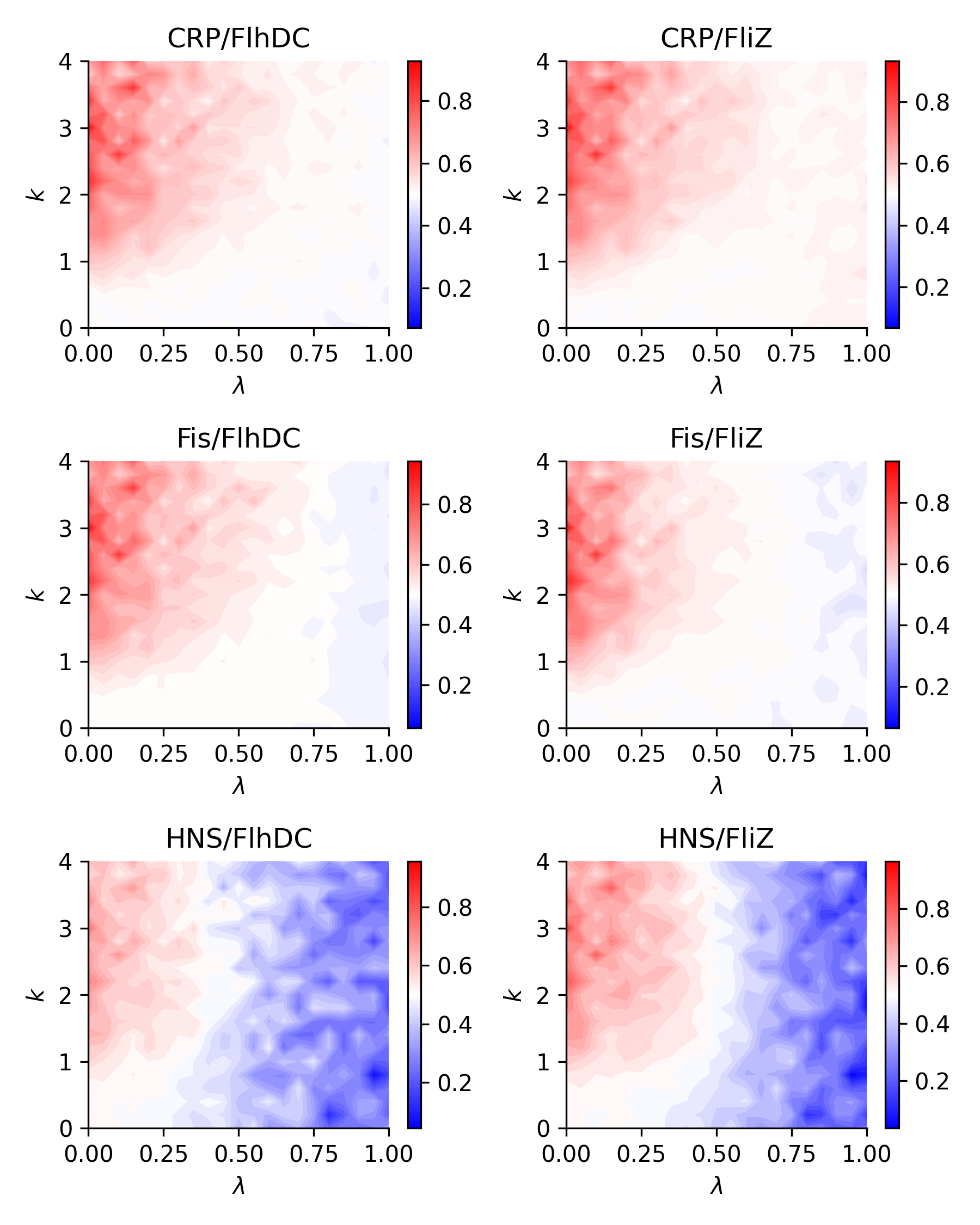

**Supplementary Figure 7**: *Fraction of global and local mutations during adaptation in variable environments for each KD pair between a local and a global regulator.*

| Genotype with largest growth | Swimming distance (cm) | Growth rate (h^-1^) |
| --- | --- | --- |
| LB | 3.37 | 0.39 |
| M63 glucose | 2.5 | 0.27 |
| M63 lactose | 2.69 | 0.28 |

**Supplementary Table 1**: *Phenotypic values of the reference strain.*

| Genotype with largest growth | Genotype with second largest growth | Count (over 18) |
| --- | --- | --- |
| ΔG + ΔL | ΔG | 12 |
| ΔL | ΔG | 3 |
| ΔL | RS | 1 |
| ΔL | ΔG + ΔL | 2 |

**Supplementary Table 2**: *Ranking of genotypes for growth over all 6 pairs in the 3 media.*
